# *In vivo* administration of anti-HIV hyper-immune eggs induces anti-anti-idiotypic antibodies against HIV gp120 peptides (fragments 308-331 and 421-438) and against HIV gp41 peptide (fragment 579-601) which have demonstrable protective capacity

**DOI:** 10.1101/2021.05.02.442298

**Authors:** Angel Justiz-Vaillant, Belkis Ferrer-Cosme, Monica Fisher Smikle, Oliver Pérez

## Abstract

Isolation of antibodies from the egg yolk of chickens is of particular interest as a source of specific antibodies for oral administration to prevent infections and use them as immunodiagnostic reagents. The use of birds in antibody production results in a reduction in the use of laboratory animals. Immunized chickens produce larger quantities of antibodies (2000 mg IgY/month) than rodents (200 mg IgG/month) in the laboratory. According to Jerne’s network theory, it is possible to produce an antibody against the antigen-binding site of another antibody. This study assessed the hypothesis that immunization with viral peptides (immunogens) could provide a potent immune response that could be evaluated in chicken eggs. Human immunodeficiency virus 1(HIV-1) is used as an immunogen. The second hypothesis was that an orally administered antibody stimulates the production of a complementary antibody, the so-called anti-idiotypic antibody, which can potentially be therapeutical. This study reports and analyzes the use of eggs as therapeutic agents. We wanted to test the hypothesis that feeding chicks with hyperimmune eggs stimulates the production of anti-anti-idiotypic antibodies that neutralize the original HIV antigen fragments 308-331 or 421-438 of gp120 or fragment 579-601 of gp41. Future research could entail an anti-idiotype strategy for prophylactic vaccines. It is vital to note that it may need an anti-idiotype response to prime immunity against an HIV viral epitope, which may be used as a secondary element. The use of anti-idiotype immune responses in infected individuals may shift the balance of the immune system, allowing the organism to manage HIV infection. Therefore, it may be an avenue for immunotherapy to improve the fight against HIV infections. However, more studies and clinical trials are required to demonstrate similar human immune responses as observed in birds.

## Introduction

The species often chosen for antisera production are rabbits or other mammals. There has been a growth in the use of hens for antibody production. Immunoglobulin (Ig)-Y is the primary antibody produced by birds and has advantages over mammalian IgG. There is no activation of the mammalian complement system, no cross-reactivity with human anti-mouse antibody (HAMA), rheumatoid factors, or human blood group antigens (lack of heteroagglutinins) [1].

According to Jerne’s network theory, antibodies contain in their variable region a representation of the ‘universe’ of antigenic structures, the idiotype. It is possible to produce antibodies against antigen-binding sites of antibodies directed to the antigen [2,3]. The production of a humoral and mucosal immune response in rabbits fed daily doses of the MOPC-315 murine IgA antibody, supporting the hypothesis that human exposure to xenogeneic antibodies, most commonly bovine milk immunoglobulins, may induce the production of anti-idiotypic antibodies [4].

We wanted to support the same hypothesis by demonstrating the animals’ capacity to orally feed hyperimmune eggs to induce systemic immune responses against the same idiotype. The study’s first set was to show that chickens immunized with the HIVgp120 peptide produced specific anti-HIV antibodies. The second set of studies demonstrated that cats fed anti-HIV hyperimmune eggs could develop antibodies that can inhibit egg yolk anti-HIV antibody binding to the HIV gp120 (fragment 254-274) peptide (original antigen), showing that the anti-HIV antibody raised in cats after feeding was anti-anti-idiotypic [3].

The main goal of this study was to raise chicken antibodies against a crucial public health microorganism, human immunodeficiency virus 1. Immunogens were prepared using this microorganism. The chickens were vaccinated intramuscularly. After booster immunization, most eggs were collected and assessed for the presence of specific antibodies. The most important result was the production of many anti-HIV antibodies in chicken eggs, with a high quantity of antibodies that inhibit the binding of Ab1 antibodies to the original antigen. Enzyme-linked immunosorbent assays were used to detect the presence of anti-HIV antibodies.

## Materials and methods

Ethical approval was granted by the Campus Ethics Committee of the University of the West Indies, Mona Campus. The laboratory work described was conducted under EU Directive 2010/63/EU for experiments.

A higher confidence level indicates a larger sample size. Power is the probability of statistically significant evidence of disparity among the groups, given the population’s variation. A higher power requires a larger sample size. A previously published HIV vaccine study included a small number of animals [4]. This new experiment documented here aims to describe earlier findings of the development of specific antibodies using oral hyperimmune eggs as feeding. This was a pilot study of an oral vaccine against HIV, using cat-fed chicken eggs as a therapeutic option [4].

The chicken house is made of wood and recreates the natural habitat of chickens. It has a black shade around the house that provides 70% shade. The area to the sides and front of the chicken house was weeded and cleaned for esthetics to minimize bacterial growth. The shaded house helped reduce the admission of sun rays, which, coupled with the zinc roof, would increase the chicken house’s temperature, which is undesirable for the birds. The chickens were housed in separate cages. Egg collection was performed daily, including on weekends.

The chicken house material consisted of a Hypo layer feed, 1.5m gauged mesh, 0.25 gauge expanding metal, and push broom shower. Other elements include water buckets, bleach (sodium hypochlorite), hydrogen peroxide (5%), egg trays, industrial gloves, sterile latex gloves, sterile applicator sticks, 27G1/2 size syringe, feed tray, and water trays.

The chickens were tested and observed for four months or another specified period. No vitamins or medications of any sort were administered to the birds under the project. The animal house was washed out with cleaning products twice daily, and fecal content was removed in keeping with a sound biosecurity practice. The chickens were watched daily for daily well-being and health and supervised by a veterinary technician. Fresh water and corn were provided to birds under free demand.

### Immunogenicity studies of experimental HIV vaccine candidates in chickens

All grade reagents used in this study were commercially available. They were purchased from Sigma-Aldrich. The study was repeated three times and similar results were obtained. The HIV immunogens used in these experiments were keyhole limpet hemocyanin (KLH) conjugated to the following HIV peptides:

1. Fragments 308-331 and 421-438 of the HIV gp120 [5]
2. Fragments 579-601 of the HIV gp 41 [5].

This peptide, derived from the V3 loop of HIV gp120, completely blocks the fusion inhibition activity of antisera to recombinant human HIV proteins and inhibits the infection of primary human T cells by HIV-1 as syncytium formation.

The second peptide, 421-438, belongs to the CD4 binding site of HIV gp120. We also used fragments 579-601 from the N-terminal heptad repeat (NHR) and loop region (loop) of HIV gp 41 (fragment 579-601) [2].

The amino acid sequences of HIV peptides and their references are cited below [5]. HIV gp 41 (fragment 579-601): Arg-lle-Leu-Ala-Val-Glu-Arg-Tyr-Leu-Lys-Asp-Gln-Gln-Leu-Leu-Gly lle-Trp-Gly Cys-Ser-Gly Lys [6].

HIV gp120 (308-331): Asn-Asn-Thr-Arg-Lys-Ser-lle-Arg-lle-Gln-Arg-Gly-Pro-Gly-Arg-Ala-Phe-Val-Thr-lle-Gly-Lys-lle-Gly [7].

HIV gp120 (421-438): Lys-Gln-Phe-lle-Asn-Met-Trp-Gln-Glu-Val-Gly-Lys-Ala-Met-Tyr-Ala-Pro-Pro [8].

### Dimerization of HIV peptides and preparation of HIV immunogens

The C-terminal cysteine was added to the amino acid sequences of HIV peptides (fragment 579-601 of HIV gp 41 and fragments 308-331 and 421-438 of HIV gp120. These peptide fragments are dimerized via cysteine oxidation in dimethyl sulfoxide [9]. Each HIV peptide was dissolved in 5% acetic acid to a final concentration of 5.1 mg/mL. The pH of the medium was adjusted to 6 with 1 M (NH4)2CO3, and dimethyl sulfoxide was added to 20% of the final volume, and after four h at room temperature (RT), the solute was extracted. Subsequently, the peptide was dissolved in 3 mL of 5% trifluoroacetic acid and precipitated with 35 mL of cold ether. The precipitate was dialyzed against 1.2 L of deionized water, pH 7 at 4°C overnight [2].

Then, 1 mg of keyhole limpet hemocyanin (KLH) was diluted in 2.1 ml 0.1 M borate buffer (1.24 g boric acid, 1.90 g sodium tetraborate, pH 10, in 500 mL deionized water). In a 20 ml glass tube, with gentle stirring, 1.1 µmol of each HIV synthetic peptide (with C-terminal cysteine added) and 0.22 milliliters 0.3% glutaraldehyde solution (ACS reagent grade, pH 5.5, Sigma-Aldrich) at RT were slowly mixed and left to stand for 1.50 hrs. A yellow coloration was observed, indicating that the conjugation process was successful. To blocking the excess of glutaraldehyde, 0.26 ml of 1 M glycine (Sigma-Aldrich) was added to block the excess glutaraldehyde, and the mixture was left for 32 min at RT. Each HIV-hemocyanin conjugate was then dialyzed against 1.3 liters 0.1 M of borate buffer, pH 8.4 through the night at 4°C. The same buffer was then used to dialyze the preparations for 8 h at 4°C. The dialysates were stored at 4°C until further use. 2.

### Chicken immunization

Nine healthy brown Leghorn layer hens (three per HIV immunogen), aged seven months, were immunized intramuscularly (IM) at multiple sites on the breast with a specific KLH-conjugated HIV peptide vaccine (KLH-conjugated HIV peptides were HIVgp41 fragment 579-601, HIVgp120 fragment 308-331, or HIVgp120 fragment 421-438). Chickens were vaccinated on day 0, with 0.5 mg/mL of the immunogens in 0.5 ml of complete Freund’s adjuvant (Sigma-Aldrich), and on days 14, 28, and 45 after the first immunization, hens received booster doses of 0.25 mg/mL of the immunogen in 0.5 mL incomplete Freund’s adjuvant. Eggs were collected daily before and after the immunization.

Additionally, nine healthy brown Leghorn chickens were fed hyper-immune eggs for 45 days (three birds for each candidate vaccine: KLH conjugated to HIV peptides, HIVgp41 fragment 579-601, HIVgp120 fragment 308-331, or HIVgp120 fragment 421-438). Likewise, three healthy brown Leghorn chickens were fed non-hyper-immune eggs for 45 days, and their response to each peptide was measured. The procedure was blinded to the procedure. The scientist did not intervene in the feeding of the birds nor knew none of the groups.

### Feeding of chicks

Feeding of chicks with HIV hyper-immune eggs with a titer of 1:15000 was fed to 15) chicks aged zero and divided into three groups fed for one month. The first group of five chicks was supplied with hyper-immune eggs against fragment 579-601 of the HIV gp 41 (fragment 579-601) peptide and corn. Likewise, the second group of five chicks was fed a hyper-immune egg against segments 308-331 of the HIV gp120 peptide and corn. The third group of five chicks was supplied with a hyper-immune egg against fragments 421-438 of the HIV gp120 peptide and corn. An additional ten chicks were divided into two groups: a group of five chicks that received non-hyperimmune HIV eggs and corn, and the last group of five chicks was fed with only corn. The feeding schedule was a blinding up process, the scientist did not take part in this process, and they did not know that each group was being treated. At the end of the feeding schedule, blood samples were obtained from all the chicks and investigated using an inhibition ELISA [5].

The water-soluble fraction (WSF) contains an elevated IgY concentration, which is separated from the lipid content by the partial application of the Polson methodology [9], which uses only chloroform, as follows: Despues de : van minúsculas y termina con ; y ultimo se une con and

1. wash the egg with warm water and crack the eggshell carefully;
2. The egg yolk was manually separated from the egg white, and the egg content was placed on tissue paper to remove as much egg white as possible.
3. The egg yolk was washed four times with 100 mL of PBS (pH 7.4) (Sigma-Aldrich).
4. break the egg yolk membrane and pour it into a 50 mL tube. The egg yolk was diluted 1:5 in PBS (pH 7.4), and an equal volume of chloroform (ACS reagent grade, Sigma-Aldrich) was added.
5. The mixture was shaken and vortexed, and the tube containing the mixture was rolled on a rolling mixer for 15 min.
6. centrifuge the mixture for 15 min (2000×g at 18°C); and
7. decant the supernatant, which contains the diluted egg yolk plasma WSF rich in IgY (25 mg/mL), which is approximately four times the IgY concentration present in the serum (6 mg/mL).

### Indirect enzyme-linked immunosorbent assay

#### Preparation of ELISA reagents

The coating buffer was prepared by adding 3.7 g Sodium Bicarbonate (NaHCO3) and 0.64 g Sodium Carbonate (Na2CO3) in 1 L of distilled water. Phosphate buffered-saline Tween-20 (10% PBS-Tween 20, pH 7.2) was made by dissolving the following reagents 0.2 g of KCl, 8 g of NaCl, 1.45 g of Na2HPO4, 0.25 g of KH2PO4, and 2 mL of tween-20 in 800 mL of distilled water. The pH was adjusted to 7.2, and distilled water was added to adjust the volume to 1 L, which was then sterilized by autoclaving. The blocking solution was prepared by mixing 0.1 g KCl, 0.1 g K3PO4, 1.16 g Na2HPO4, and 4.0 g NaCl in 500 mL distilled water, pH 7.4. Fifteen grams of non-fat dry milk was added to complete the preparation of this solution, and 15 g of non-fat dry milk was added. The sample/conjugate diluent was prepared by adding 15 g of non-fat dry milk and 2.5 mL of 10% Tween 20 to 500 mL of PBS [5].

#### ELISA for anti-HIV peptide antibodies

The 96-well polystyrene microplates (U-shaped bottom, Sigma-Aldrich) were coated with 100 ng of fragment 579-601 of HIV gp 41, fragments 308-331, or fragment 421-438 from HIV gp120 in coating buffer for 1 h at 37°C. Each microplate was washed four times with 10% PBS-Tween 20, and the blocking solution (3% non-fat milk in PBS) was added to each well (51 µL). The microplates were incubated for 1.30 h at RT. The microplates were washed as previously described. Fifty microliters of WSF diluted 1:50 with the sample diluent was added to the wells. Each microplate was then incubated for 1 h at RT and washed four times, as previously described. Then, 50 µL of horseradish peroxidase-labeled anti-IgY conjugate (Sigma-Aldrich) diluted 1:30,000 was poured into each well. The microtiter plates were incubated again for 1h at RT and washed four times. A volume of 50 µL tetramethylbenzidine (TMB, Sigma-Aldrich) was added. After further incubation for 16 min in the dark, the reaction was stopped with a solution of 3M HCl, and each microplate was read in a microplate reader at 450 nm. The cut-off value was assessed based on the mean optical density (OD) of the negative control time of 2 [2-5]. The cut-off points of ELISAs for the detection of the anti-HIV peptide (579-601), anti-HIV peptide (308-331), and anti-HIV peptide (421-438) were 0.42, 0.40, and 0.44, respectively.

Positive and negative controls were homemade. Four positive controls were used in each assay, prepared from egg yolk samples with the highest titers of specific anti-HIV peptide antibodies, and their OD values were between 1.20 and 1.50 at 450 nm. Four negative controls were used in each assay. They were prepared from the egg yolks of non-immunized animals, and they showed OD values of 0.170-0.20 at 450 nm. Three replicates of each WSF sample per bird, collected on day 60, were assayed for the presence of anti-HIV peptide antibodies using ELISA.

#### Inhibition immunoassays

Ninety-six-well polystyrene microplates (U-shaped bottom, Sigma-Aldrich Co, St Louis Missouri) were coated with 50 µL/well of 1 ng/µL solution of fragments 308-331 or 421-438 from HIV gp120 peptides or fragment 579-601 of HIV gp41 (Sigma-Aldrich) in carbonate-bicarbonate buffer (pH 9.6; Sigma-Aldrich) for 4 h at 37°C. The microplates were then washed four times with PBS Tween-20 and blocked with 3% non-fat milk in PBS, 25 µL/well, 1h, RT). Triplicates of serial doubling dilutions of chick sera diluted in PBS 3% non-fat milk (pH 7.4) were added to the microplate, which was then incubated for 90 min at 37°C. The microplates were washed four times with 150 µL of PBS-Tween 20, and 25 µL of a WSF with anti-HIV gp120 (fragments 308-331 or 421-438) antibody titer of 1:5000 or anti-HIV gp41 (fragments 579-601) with an antibody titer of 1:1000 were added to each well. Following incubation for 90 min at 37°C, the microplates were washed 4X and 25 µL of a 1:30,000 dilution of rabbit anti-chicken IgY-HRP was added to each well. The microplates were then incubated at 37°C for 1 h before final washing (PBS-Tween 20, 4 times), and 25 µL of TMB was added before incubation in the dark for 15 min. The reaction was stopped using 3M H2SO4. Microplates were read at 450 nm wavelength using a microplate reader. The percentage inhibition (%I) was calculated using the following formula: The intra-assay coefficient of variation was then calculated.

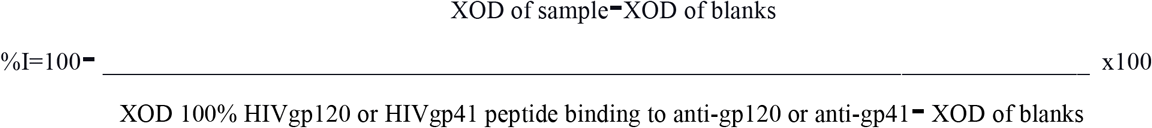

## Results and Discussion

Table 1 shows the water-soluble fractions (WSF) of egg yolks collected at zero and 60-days post-immunization were assayed for the presence of anti-HIV peptide antibodies using ELISA (six samples in total were assayed and evaluated three times). These immunogenicity results showed that HIV peptide vaccines effectively produced strong anti-HIV immune responses in immunized brown Leghorn layer hens. There was a statistically significant difference (P<0.01) that explains the notable degree of difference (related to the anti-HIV antibody levels in the egg yolks) between pre-immunized and post-immunized birds in the three experimental vaccines [2]. These immunogenicity results showed that HIV peptide vaccines effectively produced a strong anti-HIV immune response in immunized brown Leghorn layer hens.

**Table 1.** Results of immunogenicity studies of experimental HIV vaccine in brown Leghorn layer hens Mean optical density (XOD), Pre- (day 0) and post-immunization (d 45) titers, SD, standard deviation.

| Candidate vaccines | XOD (SD), day 0 (Pre-immunized birds) | XOD (SD), 45 days post-immunized birds | P-value |
| --- | --- | --- | --- |
| HIV gp 41 (fragment 579-601) 3 birds | 0.170 (0.021) | 0.885 (0.044) | <0.001 |
| HIV gp120 (fragment 308-331) 3 birds | 0.156 (0.015) | 0.910 (0.023) | <0.001 |
| HIV gp120 (fragment 421-438) 3 birds | 0.188 (0.01) | 0.865 (0.037) | <0.001 |

The results showed nine healthy brown Leghorn layer hens (three birds *per* HIV candidate vaccine), aged seven months, that were immunized IM at multiple sites on the breast with three specific KLH-conjugated HIV peptide vaccines (five birds *per* candidate) on days 0, 15, 30, and 45. Eggs from the pre-immunized and post-immunized birds were collected. The WSF of each egg [9] was evaluated using an ELISA kit. The coefficient of variation was less than 10%. This suggests that the assays are reproducible (Supporting Information). We could not compare our HIV tests with a commercially prepared assay because such a study was not available at the time of this study, and it is still not known.

Table 2 shows a significant statistical significance between birds fed non-hyperimmune eggs and birds fed hyperimmune eggs 45 days post-immunization. This corroborates that hyper-immune eggs cause oral immunization and can be considered an anti-idiotypic vaccine.

**Table 2.** Results of immunogenicity studies of experimental HIV vaccine in brown Leghorn layer hens Mean optical density (XOD) and standard deviation of birds fed or not hyperimmune egg for 45 days.

| Candidate vaccines | XOD (SD), day 45 (Birds fed with non-hyper-immune eggs) | XOD (SD), 45 days (Birds fed with hyper-immune eggs) | P-value |
| --- | --- | --- | --- |
| HIV gp 41 (fragment 579-601) | 0.250 (0.023) | 0.995 (0.068) | 0.001 |
| HIV gp120 (fragment 308-331) | 0.312 (0.017) | 0.859 (0.040) | <0.001 |
| HIV gp120 (fragment 421-438) | 0.278 (0.012) | 0.773 (0.021) | <0.001 |

Table 3 shows the results of the inhibition of fragments of HIV gp120 (fragments 308-331 or 421-438) or HIV gp41 (fragment 579-601) and anti-HIV gp120 or anti-HIV gp41 reactions by anti-anti-idiotypic HIV gp120 (fragments 308-331 or 421-438) or HIV-gp41 antibodies, respectively. This inhibition was statistically significant (P <0.05).

**Table 3.** Inhibition experiments. Mean of binding inhibition percentage (X%I), and Standard deviation (SD).

| Candidate vaccines | X%I (SD) of non-fed chicks with hyper-immune eggs and corn | X%I (SD) of fed chicks with hyper-immune eggs and corn (15-days post-immunization) | p-value |
| --- | --- | --- | --- |
| HIV gp 41 (fragment 579-601) | 3.13 (0.37) | 18.05 (2.40) | 0.007 |
| HIV gp120<br>(fragment<br>308-331) | 2.41 (0.63) | 19.62 (3.13) | 0.009 |
| HIV gp120<br>(fragment<br>421-438) | 3.19 (0.51) | 13.92 (3.96) | 0.041 |

The binding inhibition assay showed that 13.9%–20.3% inhibition of the binding of avian anti-HIV gp 41 (fragment 579-601) or anti-HIV-gp120 antibodies (Ab-1) to immobilized HIVgp41, HIVgp120 peptide by anti-HIV gp 41 (fragment 579-601) or anti-HIV gp120 (fragments 308-331 or 421-438) antibodies present in serum sample replicates of chicks tested, suggesting that the chicks anti-HIV gp 41 (fragment 579-601) or anti-HIV-gp120 antibodies (fragments 308-331 or 421-438) were anti-anti-idiotypic antibodies. This inhibition was not observed in the sera of chicks that were not fed with hyperimmune eggs. This confirms the hypothesis that feeding chicks with hyperimmune eggs stimulates the production of anti-anti-idiotypic antibodies that neutralize the original HIV antigen (gp120 or gp41).

Table 4 shows that chicks fed with corn and water failed to inhibit the binding or reactivity of fragments of HIV gp120 (fragments 308-331 or 421-438) or HIV gp41 (fragment 579-601) and anti-HIV gp120 (fragments 308-331 or 421-438) or anti-HIV gp41 (fragment 579-601) by anti-anti-idiotypic HIV gp120 (fragments 308-331 or 421-438) or HIV-gp41 antibodies, respectively; and again the chicks fed with hyper-immune eggs inhibited successfully the reactivity, suggesting that the anti-anti-HIV antibody against each fragment of gp120 or gp41 was able to recognized the original antigen and therefore reacting and protecting against it.

**Table 4.** Inhibition of fragments of HIV gp120 (fragments 308-331 or 421-438) or HIV gp41 (fragment 579-601) and anti-HIV gp120 (fragments 308-331 or 421-438) or anti-HIV gp41 (fragment 579-601) reaction by anti-anti-idiotypic HIV gp120 (fragments 308-331 or 421-438) or HIV-gp41 antibodies, respectively. Mean of binding inhibition percentage (X%I), and Standard deviation (SD).

| Candidate vaccines | X%I (SD) of non-fed chicks with hyper-immune eggs (given corn and water) | X%I (SD) of fed chicks with hyper-immune eggs and corn (30-days post-immunization) | p-value |
| --- | --- | --- | --- |
| HIV gp 41 (fragment 579-601) 3 birds | 3.13 (0.37) | 20.08 (3.23) | 0.011 |
| HIV gp120 (fragment 308-331) 3 birds | 2.41 (0.63) | 18.55 (2.47) | 0.005 |
| HIV gp120 (fragment 421-438) 3 birds | 3.19 (0.51) | 16.79 (3.85) | 0.024 |

**Table 5.** Inhibition of fragments of HIV gp120 (fragments 308-331 or 421-438) or HIV gp41 (fragment 579-601) and anti-HIV gp120 (fragments 308-331 or 421-438) or anti-HIV gp41 (fragment 579-601) reaction by anti-anti-idiotypic HIV gp120 (fragments 308-331 or 421-438) or HIV-gp41 antibodies, respectively. Mean of binding inhibition percentage (X%I), and Standard deviation (SD).

| Candidate vaccines | X%I (SD) of fed chicks with hyper-immune eggs (15-days post-immunization) | X%I (SD) of fed chicks with hyper-immune eggs (30-days post-immunization) | t-value | p-value |
| --- | --- | --- | --- | --- |
| HIV gp 41 (fragment 579-601) | 18.05 (2.40) | 20.08 (3.23) | -1.24 | 24.71% |
| HIV gp120 (fragment 308-331) | 19.62 (3.13) | 18.55 (2.47) | 0.66 | 52.66% |
| HIV gp120 (fragment 421-438) | 13.92 (3.96) | 16.79 (3.85) | -1.27 | 23.19% |

These experiments did not reject the null hypothesis, indicating that there was no statistically significant difference between the variables analyzed. Birds fed hyper-immune eggs 15 days or 30 days post-immunization were able to induce the same stimulated immune response against the original antigen: HIV gp120 (fragments 308-331 or 421-438) or HIV gp41 (fragment 579-601).

It was shown that feeding chicks with anti-idiotypic HIV gp41 (fragment 579-601) or gp120 peptide antibodies induced anti-peptide antibodies (Ab-3), which also recognized a recombinant HIV gp41 (fragment 579-601) or HIV gp120 (fragments 308-331 or 421-438), and also inhibited its interaction with anti-HIV-gp41 or anti-HIV-gp120 antibodies (Ab1). This study provides evidence that is supported by a previous study in which intramuscular immunization of BALB/c mice with anti-idiotypic HIV gp160 peptide antibodies induced anti-peptide antibodies (Ab-3), which also recognized a recombinant HIV gp160 preparation, and inhibited its interaction with anti-HIV-gp160 antibodies (Ab1) [11]. Anti-idiotypic vaccines in HIV/AIDS, using a monoclonal antibody (mAb 13B8.2) directed against the CDR3-homologous CD4/D1 region has previously been reported in the literature [4,12]. Our results are also supported by a study on the development of an idiotype-anti-idiotype network of antibodies to bovine serum albumin (BSA) in eggs from chickens immunized with BSA [13]. It has also been reported that HIV-1 gp160-specific secretory IgA was detected in the saliva of all rabbits orally immunized with HIV-immunosomes, neutralizing HIV infectivity in vitro [14]. Our study showed a statistically significant percentage inhibition. Oral immunizations had a p-value < 0.05, which indicated the presence of anti-anti-idiotypic antibodies (Ab3) against HIV gp120 (fragments 308-331 or 421-438) and HIV gp 41 (fragment 579-601) fragments, which inhibited the formation of complexes between HIV gp120 (fragments 308-331 or 421-438) peptide or HIV gp 41 (fragment 579-601) and anti-HIVgp120 or anti-HIV gp 41 (fragment 579-601) idiotypic antibodies, respectively.

Boudet *et al. (*1992) reported that one of the well-characterized motifs mapped to a loop within the third hypervariable region (V3) of the exterior envelope glycoprotein gp120 at amino acid positions 308–331 and is referred to as the principal neutralizing determinant (PND). The sequence of this V3 loop raises the immunogenicity and diversity of the antibody response to PND. We show here that this neutralization-related motif is highly immunogenic in HIV-positive subjects and experimentally immunized primates and rodents subjected to various anti-HIV immunization regimens [15]. Our study showed that this peptide could generate a robust immune response in chickens, consistent with previous research by Boudet et al. The immunogenicity study was statistically significant (p <0.05), suggesting that it could be a good immunogen for a candidate vaccine.

Bell *et al*. (1992) reported the use of a series of overlapping synthetic peptides derived from a conserved region of the envelope gp41 (aa 572-613). They identified an immunodominant 12-mer peptide sequence, gp41(8) (aa 593-604), which consistently elicited both T cell blastogenic and antibody responses in asymptomatic HIV-seropositive individuals but not in ARC and AIDS patients [16]. Linear regression analysis showed that in asymptomatic persons, there was a strong positive correlation (P less than 0.0005) between the absolute CD8+ T cell and the magnitude of blastogenic responses to gp41(8) (aa 593-604) [16]. These experiments suggest that the involved peptides have a high immunogenicity capacity, as suggested by our study. Therefore, the fragment HIV gp 41 (fragment 579-601) (572-613) is a promising vaccine candidate, as shown in our study in immunogenicity studies and binding inhibition assays.

The first (B138) is linear and spans the envelope residues 421-438; the second (1005/45) encompasses amino acids 418-445 and is cyclized by a disulfide bond joining its terminal cysteines. Both species have been shown to inhibit syncytia formation in a conventional bioassay, with B138 being the most efficient. Both peptides elicit high titers of anti-peptide antibodies in immunized mice, rabbits, and goats, with titers exceeding 1:10 in many cases [8]. Our study does not show that anti-idiotypic antibodies inhibit syncytia formation, because this was not our aim. We are not aware of the literature on other immunogenicity studies performed with the HIV gp120 (fragments 308-331 or 421-438)peptide fragment 421-438. Our study makes an original contribution to the literature.

According to Jerne’s network theory, antibodies have in their variable regions an image of the ‘universe’ of antigenic arrangements, the idiotype (Id). Therefore, antibodies can be recognized as antigens by the immune system under normal circumstances, and these interactions are called Id-anti-Id reactions, which can be used to manipulate the immune system [17].

Egg yolk is a critical and alternative source of polyclonal antibodies. Of course, they present some advantages over mammalian antibodies in terms of animal welfare, productivity, and specificity [1]. Moreover, because of its phylogenetic distance and structural molecular features, IgY is more suitable for treatment and diagnostic purposes than mammals because of its lack of reactivity with the human complement system and B cell repertoire [1]. However, IgY has a greater avidity for conserved mammalian polypeptides [18,19]. The main antibody (IgY) present in avian blood is transmitted to the offspring and stored in egg yolks, facilitating the non-invasive harvesting of many antibodies [20,21]. Indeed, chicken IgY antibodies can be used therapeutically and in rapid diagnostic tests [22-25].

Chicken IgY provides a more ethical and humane approach to therapeutic and diagnostic methods that use these polyclonal antibodies. It has a lower cost of return than mammalian antibodies [26], with the ability to produce more antibodies with fewer animals involved. It can translate to lower prices for diagnostic tests or therapeutic methods that utilize chicken IgY. Diagnostically, IgY does not cross-react with Fc receptors of rheumatoid factors, HAMA, human blood group antigens, or activates the complement system. Thus, there are fewer false-positive results when using IgY in immunological assays, making it more specific to detect several antigen classes [1,26].

Therapeutically, IgY has the potential to be used in the treatment of many infections. These include infectious diseases caused by *Helicobacter pylori* [27], enterotoxigenic *E. coli*, and *Salmonella* spp. [26]. Other problems include acute and chronic pharyngitis [28] and infantile rotavirus enteritis [29]. The prevention of oral colonization by *Streptococcus mutans* [30] and the potential for greater immunotherapy (such as for HIV) make IgY critical in modern medical treatment. Due to IgY properties, less toxicity and side effects, and fewer allergic reactions make it more suitable than previously considered immunotherapies involving mammalian antibodies. Its therapeutic potential makes it a necessary treatment that requires more consideration, especially with bacterial resistance to existing antibiotics and the emergence of novel strains of viruses [31]. Egg yolk antibodies were also obtained. Egg white immunoglobulin with specificity to staphylococcal protein A was detected in the immunized hens. It inhibits S. aureus growth in vitro [32]. A sandwich ELISA showed anti-SpA antibodies in hens immunized with titers of at least 5-folds compared to pre-immunized hens ten days post-immunization [32]. Therefore, the whole egg could be considered as a drug to combat microorganisms. In vitro experiments support the hypothesis that the idiotypic-anti-idiotypic network plays a role in immunoprotection, which deserves further study. The authors suggest that eggs from chickens immunized with specific immunogens may be considered a special type of oral anti-idiotypic vaccine. It was shown by development of anti-anti-idiotypic antibodies that were developed against various epitopes [33].

## Conclusions

Future research could entail an anti-idiotype strategy for prophylactic vaccines. It is vital to note that it may need an anti-idiotype response to prime immunity against an HIV viral epitope, which may be used as a secondary element. The use of anti-idiotype immune responses in infected individuals may shift the balance of the immune system, allowing the organism to manage HIV infection. We hypothesized that feeding chicks with hyperimmune eggs would stimulate the production of anti-anti-idiotypic antibodies that neutralize the original HIV antigen (gp120 or gp41). Therefore, it may be an avenue for immunotherapy to improve the fight against HIV infections. However, more studies and clinical trials are required to demonstrate similar human immune responses as observed in birds.

